# Longitudinal Reorganization of Large-Scale Functional Networks in SCA7

**DOI:** 10.64898/2026.07.28.741349

**Authors:** Alireza Aleali, Luis Beltran-Parrazal, Juan Fernandez-Ruiz, Carlos R. Hernandez-Castillo

**Affiliations:** Department of Medical Neuroscience, Dalhousie University, Halifax, Nova Scotia, Canada; Faculty of Computer Science, Dalhousie University, Halifax, Canada; Instituto de Investigaciones Cerebrales, Universidad Veracruzana, Xalapa, Veracruz, México; Instituto de Neuroetología, Universidad Veracruzana, Xalapa, Veracruz, México; Departamento de Fisiología, Facultad de Medicina, Universidad Nacional Autónoma de México, Ciudad de Mexico, México

**Author notes:** Correspondence to: Prof. Carlos R. Hernandez-Castillo, Ph.D., Room 438 Goldberg Computer Science Building, Dalhousie University 6050 University Avenue, Halifax, Nova Scotia, Canada B3H 4R2.

**Keywords:** Spinocerebellar ataxia type 7, Resting-state functional MRI, Cerebellar network, Longitudinal neuroimaging, Machine learning classification

## Abstract

Spinocerebellar ataxia type 7 (SCA7) is a rare neurodegenerative disorder characterized by progressive cerebellar ataxia and visual impairment. We investigated longitudinal changes in resting-state functional connectivity and their clinical associations. Resting-state functional MRI was acquired from 16 individuals with SCA7 and 16 age- and sex-matched healthy controls across three visits over 24 months. Network-to-network functional connectivity was quantified, and machine-learning models were trained using functional connectivity features. SCA7 showed lower MoCA (p = 0.045) and MMSE (p = 0.025) scores and progressive worsening of ataxia (SARA, p < 0.001). Significant Group × Visit interactions were observed for Visual–Default Mode (p = 0.012) and Somatomotor–Cerebellar Dorsal Attention connectivity (p = 0.038). Functional connectivity abnormalities involved cortical, cortico-cerebellar, and cerebellar networks that became more widespread at the final follow-up assessment, with the visual network emerging as the most consistently affected system across analyses. Functional connectivity abnormalities were associated with cognitive performance (MMSE: r = −0.62, p = 0.01) and disease severity (SARA: r = 0.589, p = 0.016). Functional connectivity features accurately classified SCA7 and healthy controls (accuracy = 96.4%, F1 = 0.969). These findings support resting-state functional connectivity as a candidate biomarker warranting further validation in larger, independent cohorts.

## 1. Introduction

Spinocerebellar ataxia type 7 (SCA7) is a rare autosomal dominant neurodegenerative disorder caused by a pathogenic expansion of cytosine–adenine–guanine (CAG) repeats in the *ATXN7* gene (David et al., 1997; Gouw, 1998; Holmberg, 1998; Hugosson et al., 2009; Johansson, 1998; Tang et al., 2000; Tezenas Du Montcel et al., 2014). Clinically, SCA7 is characterized by progressive cerebellar ataxia and cone–rod retinal degeneration with progressive macular involvement, distinguishing it from other spinocerebellar ataxias by its characteristic association with progressive visual impairment (Aleman et al., 2002; Campos-Romo et al., 2018; Michalik et al., 2004; Niu et al., 2018). Depending on the age at onset and disease stage, visual symptoms may precede, accompany, or follow the development of motor manifestations and can progress to severe vision loss (David, 1998; McLaughlin & Dryja, 2002; on behalf of Rede Neurogenetica et al., 2019; Rüb et al., 2013). In addition to gait and limb incoordination, affected individuals may develop dysarthria, dysphagia, and oculomotor abnormalities as the disease advances (Goswami et al., 2022; Horton et al., 2013a; Peng et al., 2023; Rosini et al., 2020; Seidel et al., 2012). The clinical presentation is heterogeneous, with earlier onset and more rapid disease progression generally associated with larger CAG repeat expansions (Karam & Trottier, 2018; Yamada et al., 2007). Given its progressive course and its substantial impact on both motor and visual functioning, SCA7 represents a considerable source of disability and reduced quality of life for affected individuals and their families (Park et al., 2020; Triangto et al., 2022).

Although SCA7 has traditionally been viewed as a disorder characterized by cerebellar degeneration, accumulating evidence suggests that its neuropathological involvement extends beyond the cerebellum (Garden et al., 2002; Niewiadomska-Cimicka & Trottier, 2019). Neuropathological investigations have documented neuronal loss affecting the cerebellar cortex, inferior olivary complex, and brainstem nuclei, as well as degeneration involving subcortical structures such as the pallidum and substantia nigra (Rüb et al., 2013). Consistent with these observations, structural MRI studies have identified gray matter abnormalities not only in the cerebellum and pons but also in several cerebral regions, including sensorimotor, occipital, insular, and frontal cortices (Alcauter et al., 2011; Hernandez-Castillo, Galvez, et al., 2016a). Diffusion imaging studies have further demonstrated widespread white matter abnormalities affecting cerebellar peduncles and major cortical projection pathways, with alterations in motor-and visuospatial-related tracts associated with greater ataxia severity (Hernandez-Castillo, Vaca-Palomares, et al., 2016a; Parker et al., 2021). Importantly, several of these structural abnormalities have been linked to clinical measures of disease burden, and emerging longitudinal studies have documented progressive neuroanatomical changes over time. Collectively, these findings suggest that the neurodegenerative process in SCA7 involves distributed regions of the central nervous system rather than being restricted to the cerebellum alone.

Resting-state functional MRI (rs-fMRI) has emerged as a valuable approach for investigating spontaneous functional interactions among distributed brain regions and has increasingly been applied to characterize network alterations in neurodegenerative disorders. In SCA7, an early rs-fMRI study reported disrupted functional connectivity involving visual and motor systems, including altered interactions between cerebellar regions and cerebral areas implicated in sensorimotor processing (Hernandez-Castillo et al., 2013). Subsequent whole-brain analysis extended these observations by demonstrating both increased and decreased functional connectivity across cerebellar and cerebral regions, involving not only visual and motor cortices but also temporal and prefrontal areas (Hernandez-Castillo et al., 2014). Collectively, these findings suggest that functional abnormalities in SCA7 are distributed across multiple brain systems rather than being confined to regions traditionally associated with the clinical phenotype. However, because these studies were cross-sectional, they provide only a snapshot of disease-related functional alterations. Cross-sectional abnormalities do not necessarily reflect disease progression, making it unclear whether functional changes remain stable, worsen, or reorganize as the disease advances. Consequently, the longitudinal evolution and large-scale organization of functional alterations in SCA7 remain incompletely understood.

Addressing these questions may provide a more comprehensive understanding of the functional manifestations of SCA7 and their potential clinical relevance. Therefore, in the present study, we used longitudinal resting-state fMRI to characterize functional brain organization in individuals with SCA7 and healthy controls using a hierarchical analytical framework spanning multiple levels of organization, including global functional connectivity, within- and between-network interactions, and their associations with clinical characteristics. Because imaging biomarkers may ultimately facilitate disease characterization and longitudinal monitoring, we also evaluated whether these functional measures could distinguish individuals with SCA7 from healthy controls.

## 2. Methods

### 2.1. Participants

Twenty SCA7 patients with molecular diagnosis were initially enrolled in the study. However, after completing the initial testing, four of them did not participate in the follow-up 1 or 2 years later. Reasons for drop-out included the progression toward advanced stages of the disease in three patients that precluded MRI scanning, and the passing of one patient. The remaining SCA7 group consisted of 16 patients (7 females, 9 males). We also included subject-to-subject-matched healthy participants to SCA7 patients (age-, gender-, and years of schooling-matched) as a control group. This group consisted of 16 volunteers with no history of neurological injury or psychiatric diseases. Resting-state fMRI data from four healthy controls did not meet the requirements for functional connectivity analyses because of incomplete imaging data; therefore, all functional connectivity analyses were performed using 12 healthy controls. Both groups underwent longitudinal MRI acquisitions. Participants with SCA7 additionally completed clinical and cognitive assessments at each follow-up visit, whereas these assessments were obtained from healthy controls only at baseline. The mean interval between consecutive visits was 12.83 months (SD = 0.44 months). All participants were native Spanish speakers recruited from the central region of Veracruz, México. The research protocol was approved by the Research Ethics Board of the Faculty of Medicine at the Universidad Nacional Autónoma de México (Project ID: 015/2015). Written informed consent was obtained from each participant, and all procedures were conducted in accordance with the principles of the Declaration of Helsinki.

### 2.2. Clinical Assessment

Clinical assessments were performed to characterize motor, cognitive, and affective functioning. Disease severity in participants with SCA7 was evaluated using the Scale for the Assessment and Rating of Ataxia (SARA; Schmitz-Hübsch et al., 2006), an eight-item clinical instrument that assesses gait, stance, sitting, speech disturbance, finger chase, nose–finger test, fast alternating hand movements, and heel–shin slide performance. Total SARA scores range from 0 to 40, with higher scores indicating greater ataxia severity.

Global cognitive functioning was assessed using the Montreal Cognitive Assessment (MoCA; Nasreddine et al., 2005) and the Mini-Mental State Examination (MMSE; Tombaugh & McIntyre, 1992). The MoCA is a 30-point screening tool designed to evaluate multiple cognitive domains, including attention, executive functioning, memory, language, visuospatial abilities, abstraction, calculation, and orientation. The MMSE was included as an additional measure of global cognitive status and assesses orientation, attention, memory, language, and visuo-constructive abilities. Together, these measures provided complementary assessments of cognitive performance across a broad range of domains.

Additional neuropsychological assessment included semantic and phonemic verbal fluency tasks, the Rey Auditory Verbal Learning Test (RAVLT; Rey, 1983), and the Backward Digit Span test. The semantic and phonemic verbal fluency tasks evaluate lexical retrieval, semantic memory, language production, and executive functioning through measures of word generation, clustering, and switching strategies. The RAVLT assesses multiple aspects of verbal learning and episodic memory, including immediate recall, learning rate, proactive and retroactive interference, forgetting, and recognition memory. The Backward Digit Span test was used to assess auditory working memory and executive attention by requiring participants to recall sequences of digits in reverse order.

Depressive symptoms were assessed using the Center for Epidemiologic Studies Depression Scale (CES-D; Lewinsohn et al., 1997), a self-report questionnaire that evaluates the frequency of depressive symptoms experienced during the preceding week. The CES-D is widely used in both clinical and research settings and provides a quantitative measure of depressive symptom burden.

The availability of clinical measures varied across visits. SARA scores were available only for participants with SCA7. Cognitive (MoCA and MMSE) and depression (CES-D) assessments were available for healthy controls at baseline and for participants with SCA7 at baseline and first follow-up but were not available at the final follow-up visit. Consequently, analyses involving clinical measures were restricted to visits for which the corresponding assessments were available.

### 2.3. MRI Acquisition and Preprocessing

All images were acquired with a 3 Tesla General Electric MR750 Discover system at the Instituto de Neurobiología of the Universidad Nacional Autónoma de México in Juriquilla, Querétaro, México. The study consisted in the acquisition of T1-3D anatomical high-resolution images using a SPGR sequence (Spoiled Gradient Recalled), with a TE/TR 3.18/8.16 ms; FOV 256 × 256 mm2, and an acquisition and reconstruction matrix of 256 × 256, resulting in an isometric resolution of 1 × 1 × 1 mm3. MRI images were acquired at baseline and at follow-up 24 ± 0.70 months later. Image acquisition included non-replacement of MRI scanners during the longitudinal study, no relevant hardware or software changes, and no significant changes of images quality. All images were preprocessed before the analyses, including MNI orientation, denoising, and intensity inhomogeneity correction. Functional images were collected using an Echo Planar Imaging single-shot sequence with a TR of 2,000 ms, TE of 35 ms, and 250 whole-brain volumes with 34 slices. Final isometric resolution of the functional images was 3 x 3 x 4 mm without gaps. During fMRI acquisition, subjects in all groups were instructed to keep their eyes closed, to think about nothing in particular, and to stay awake. Five dummy scans were performed at the beginning of each functional acquisition to allow magnetization to reach a steady state.

The rsfMRI preprocessing included discarding the first 4 volumes, brain extraction, time shifting, motion correction, spatial smoothing (6 mm full-width at half-maximum Gaussian kernel), linear trend removal, and temporal filtering (band pass, 0.01-0.08 Hz), using FSL (FMRIB Software Library (FSL), Oxford University, Oxford, UK). Motion-related artifacts were further removed using ICA-AROMA (Pruim et al., 2015), which classifies independent components as motion- or non-motion-related based on spatial and temporal features and removes motion-related components from the functional data. Following ICA-AROMA denoising, mean global, white matter, and cerebrospinal fluid signals were extracted and regressed from the data. All structural images were warped to the Montreal Neurological Institute MNI template, using the nonlinear registration method (Smith et al., 2004). After rigid alignment of rsfMRI images to structural images, spatial normalization of rsfMRI images to the MNI template was achieved by using the transformation field acquired during the structural image registration.

### 2.4. Functional Connectivity Analysis

To investigate longitudinal alterations in large-scale functional network organization in SCA7, we quantified network-to-network functional connectivity among cortical and cerebellar functional systems. This network-level approach enabled the characterization of cortical, cortico-cerebellar, and cerebellar interactions while reducing the dimensionality of parcel-wise connectivity into biologically interpretable measures of large-scale brain organization. Functional connectivity analyses were performed using the Schaefer-200 cortical parcellation (Schaefer et al., 2018), with cortical parcels assigned to the seven canonical Yeo functional networks (Thomas Yeo et al., 2011). To characterize cortico-cerebellar and cerebellar interactions, the seven cerebellar functional systems defined by the Buckner-7 atlas (Buckner et al., 2011) were additionally incorporated.

For each participant and visit, mean BOLD time series were extracted from the 200 cortical parcels and seven cerebellar systems, yielding 207 regional time series. Pairwise Pearson correlation coefficients were calculated between all regions to generate a 207 × 207 functional connectivity matrix, and correlation coefficients were transformed to Fisher z-scores.

To provide a broader characterization of functional brain organization, additional analyses of global functional connectivity and network organization were also performed. These included whole-brain mean functional connectivity, within-network connectivity, between-network connectivity, and system segregation and are presented in the Supplementary Material.

### 2.5. Statistical Analysis

All statistical analyses were performed using R (version 4.2.2; R Foundation for Statistical Computing, Vienna, Austria) and IBM SPSS Statistics (version 29.0.2; IBM Corp., Armonk, NY, USA). Prior to analysis, the distribution of clinical and imaging variables was assessed using the Kolmogorov–Smirnov and Shapiro–Wilk tests of normality.

Longitudinal changes in functional connectivity measures were evaluated using linear mixed-effects (LME) models. Group, Visit, and their interaction (Group × Visit) were included as fixed effects, while Subject was included as a random intercept to account for repeated measurements. Age and sex were included as covariates of no interest in all models. Statistical models were specified as:

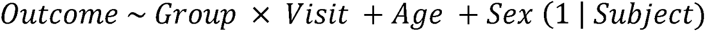

Estimated marginal means were used to perform pairwise comparisons examining between-group differences at each visit and within-group changes across visits. Associations between functional connectivity measures and clinical variables were evaluated using Pearson or Spearman correlation analyses, as determined by the results of normality testing. All clinical and behavioral measures showing significant group differences were tested for association with the significant functional connectivity findings. Correlation analyses were restricted to participants and visits for which the corresponding clinical measures were available.

To control for multiple comparisons, false discovery rate (FDR) correction was applied using the Benjamini–Hochberg procedure. FDR correction was performed separately within each family of related analyses. Statistical significance was defined as p < 0.05 after correction for multiple comparisons.

### 2.6. Machine Learning

To evaluate the potential of resting-state functional connectivity as a biomarker for distinguishing individuals with SCA7 from healthy controls, supervised machine-learning models were trained using functional connectivity features. The input features consisted of the functional connectivity measures identified as statistically significant in the preceding network-to-network analyses. This analysis was performed to assess whether large-scale functional network alterations identified through conventional statistical analyses also possessed discriminative value at the individual-subject level.

Three supervised classification algorithms were evaluated: Logistic Regression, Random Forest, and Support Vector Machine (SVM). Model performance was assessed using a nested cross-validation framework. An outer Leave-One-Out Cross-Validation (LOOCV) procedure was employed, in which a single participant was held out for testing while the remaining participants were used for model training. Within each training set, hyperparameter optimization was performed using an inner stratified k-fold cross-validation procedure. The number of folds was determined by the size of the smallest class and was limited to a maximum of three folds to maintain class representation across validation splits. Hyperparameters were optimized using a grid-search strategy within the inner cross-validation loop.

Classification performance was evaluated using accuracy and F1 score, calculated from predictions generated across all LOOCV test folds. Following model evaluation, the best-performing classifier was identified based on overall classification performance. A confusion matrix was generated for the final selected model, and feature-importance analyses were performed to identify the functional connectivity features contributing most strongly to classification performance.

## 3. Results

### 3.1 Baseline Clinical Characteristics and Longitudinal Clinical Changes in SCA7

The SCA7 and control groups were comparable in age, sex distribution, and years of education. Within the SCA7 cohort, the mean disease duration was 6.94 ± 3.33 years and the mean expanded CAG repeat length was 48.93 ± 4.44 repeats. Baseline clinical comparisons were limited to Visit 1 because clinical measures were only available for the control group at this time point. Relative to controls, participants with SCA7 demonstrated lower cognitive performance on both the MoCA and MMSE, whereas depressive symptoms assessed using the CES-D, along with semantic or phonemic verbal fluency measures, RAVLT performance, and Backward Digit Span, did not differ significantly between groups. Demographic and baseline clinical characteristics are summarized in Table 1.

**Table 1.**
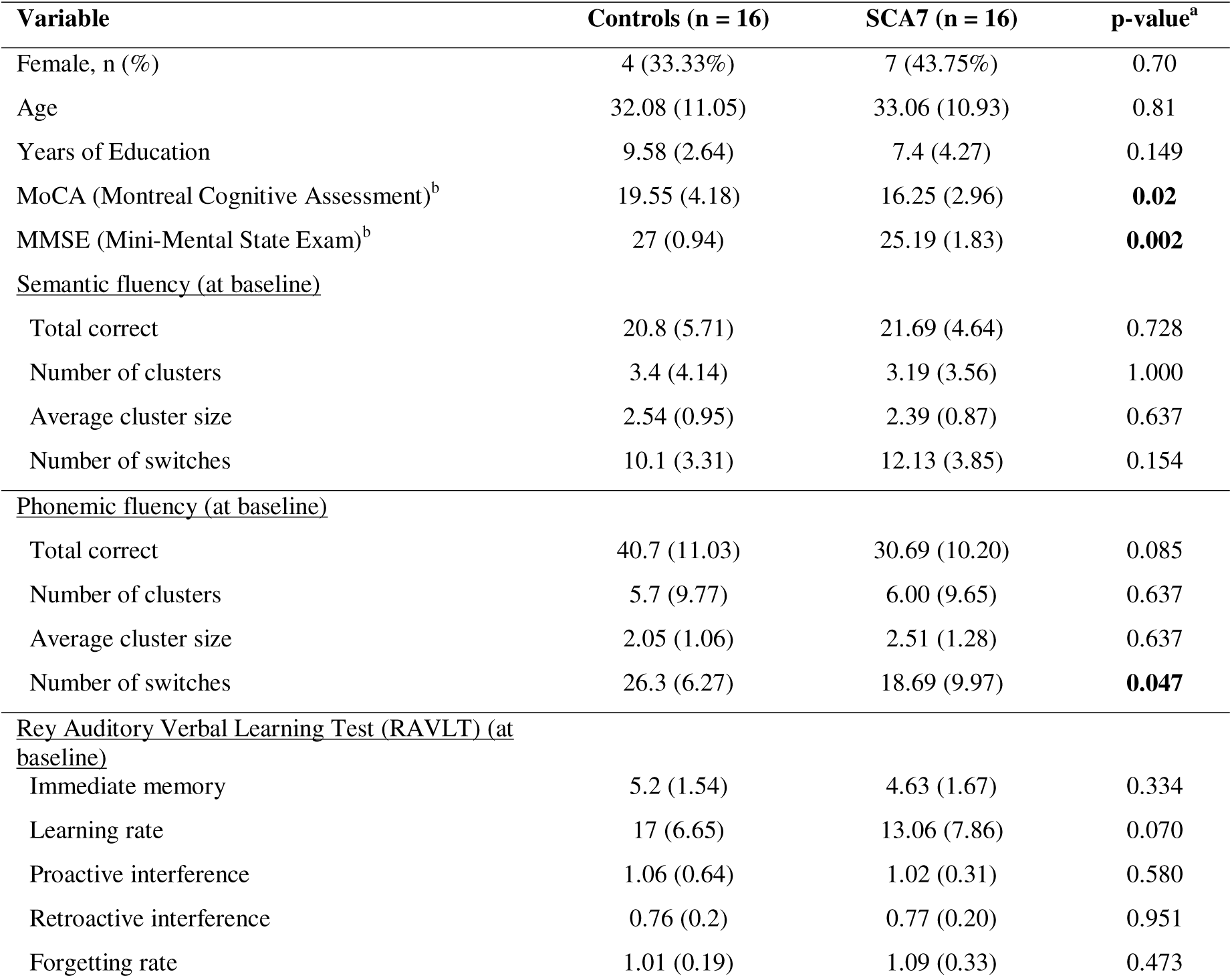
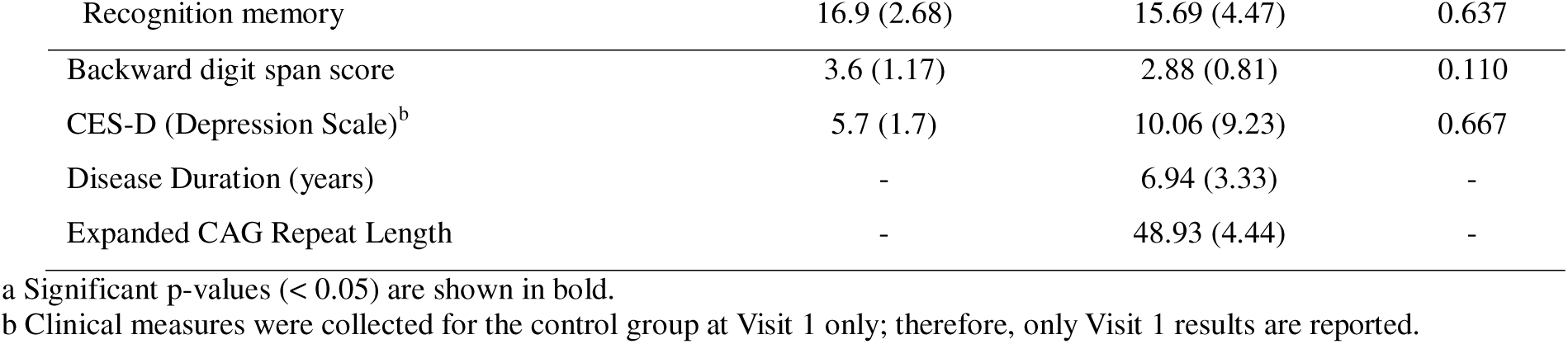
Demographic and baseline clinical characteristics of SCA7 patients and matched healthy controls. Data are presented as mean ± SD. Between-group comparisons were performed using the Mann–Whitney U test for continuous variables and the χ² test for sex distribution. SCA7, spinocerebellar ataxia type 7; CES-D, Center for Epidemiologic Studies Depression Scale; SD, standard deviation.

| Variable | Controls (n = 16) | SCA7 (n = 16) | p-value<sup>a</sup> |
| --- | --- | --- | --- |
| Female, n (%) | 4 (33.33%) | 7 (43.75%) | 0.70 |
| Age | 32.08 (11.05) | 33.06 (10.93) | 0.81 |
| Years of Education | 9.58 (2.64) | 7.4 (4.27) | 0.149 |
| MoCA (Montreal Cognitive Assessment)<sup>b</sup> | 19.55 (4.18) | 16.25 (2.96) | **0.02** |
| MMSE (Mini-Mental State Exam)<sup>b</sup> | 27 (0.94) | 25.19 (1.83) | **0.002** |
| <u>Semantic fluency (at baseline)</u> |  |  |  |
| Total correct | 20.8 (5.71) | 21.69 (4.64) | 0.728 |
| Number of clusters | 3.4 (4.14) | 3.19 (3.56) | 1.000 |
| Average cluster size | 2.54 (0.95) | 2.39 (0.87) | 0.637 |
| Number of switches | 10.1 (3.31) | 12.13 (3.85) | 0.154 |
| <u>Phonemic fluency (at baseline)</u> |  |  |  |
| Total correct | 40.7 (11.03) | 30.69 (10.20) | 0.085 |
| Number of clusters | 5.7 (9.77) | 6.00 (9.65) | 0.637 |
| Average cluster size | 2.05 (1.06) | 2.51 (1.28) | 0.637 |
| Number of switches | 26.3 (6.27) | 18.69 (9.97) | **0.047** |
| <u>Rey Auditory Verbal Learning Test (RAVLT) (at baseline)</u> |  |  |  |
| Immediate memory | 5.2 (1.54) | 4.63 (1.67) | 0.334 |
| Learning rate | 17 (6.65) | 13.06 (7.86) | 0.070 |
| Proactive interference | 1.06 (0.64) | 1.02 (0.31) | 0.580 |
| Retroactive interference | 0.76 (0.2) | 0.77 (0.20) | 0.951 |
| Forgetting rate | 1.01 (0.19) | 1.09 (0.33) | 0.473 |
| Recognition memory | 16.9 (2.68) | 15.69 (4.47) | 0.637 |
| Backward digit span score | 3.6 (1.17) | 2.88 (0.81) | 0.110 |
| CES-D (Depression Scale)<sup>b</sup> | 5.7 (1.7) | 10.06 (9.23) | 0.667 |
| Disease Duration (years) | - | 6.94 (3.33) | - |
| Expanded CAG Repeat Length | - | 48.93 (4.44) | - |

a Significant p-values (< 0.05) are shown in bold.
b Clinical measures were collected for the control group at Visit 1 only; therefore, only Visit 1 results are reported.

Longitudinal analyses were restricted to Visits 1 and 2 because clinical assessments were unavailable at the final follow-up visit. Across the observation period, participants with SCA7 exhibited a significant increase in total SARA scores, accompanied by significant changes across multiple gait, speech, upper-limb, and lower-limb coordination measures. MoCA scores also declined significantly between visits. In contrast, no significant longitudinal changes were observed for MMSE or CES-D scores. Longitudinal clinical changes within the SCA7 group are summarized in Table 2.

**Table 2.**
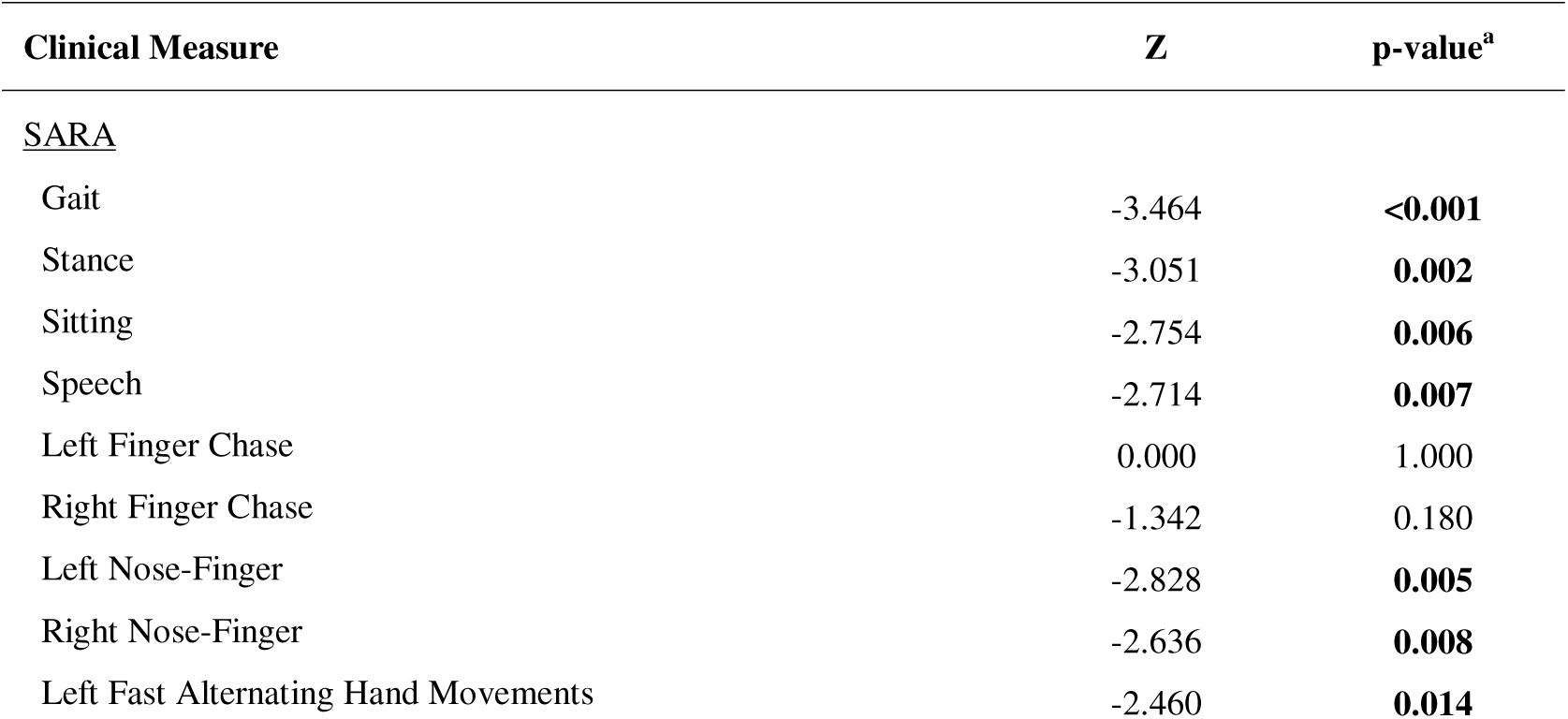
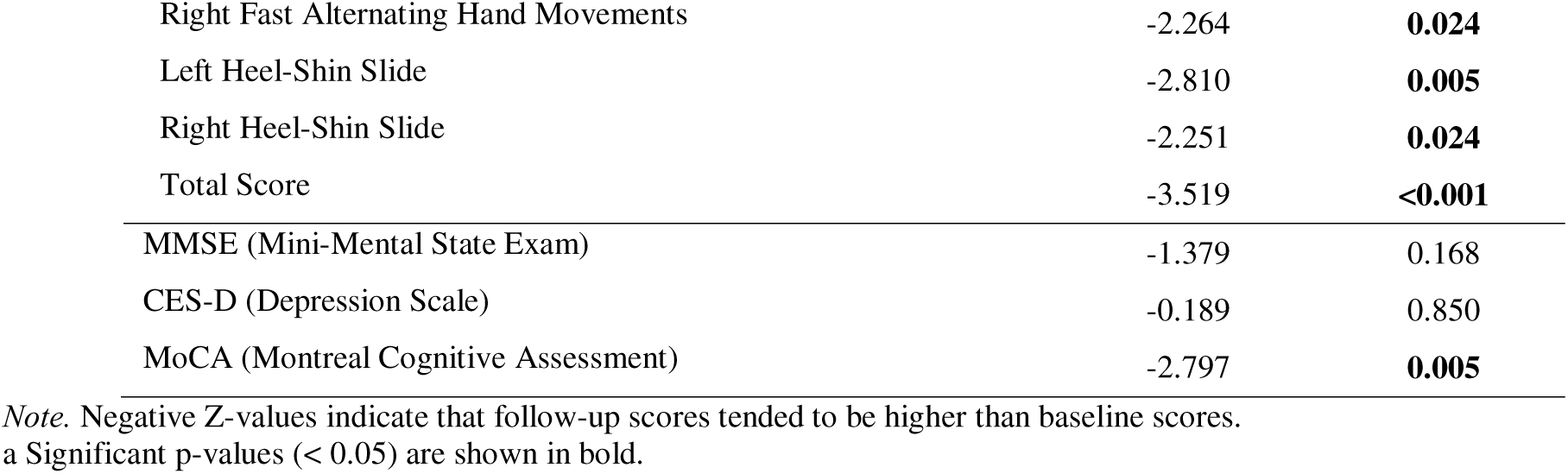
Longitudinal changes in clinical measures between Visit 1 and Visit 2 in participants with SCA7. Data are presented as mean ± SD. Longitudinal comparisons were performed using the Wilcoxon signed-rank test. Clinical assessments were unavailable at Visit 3 and were therefore excluded from longitudinal analyses. SARA, Scale for the Assessment and Rating of Ataxia; CES-D, Center for Epidemiologic Studies Depression Scale; SD, standard deviation.

### 3.2 Visual–Default and Cortico-Cerebellar Interactions Exhibited Distinct Longitudinal Alterations in SCA7

Between-system integration analyses identified two network pairs exhibiting significant Group × Visit interaction effects following FDR correction (Figure 1). Connectivity between the Cortical Visual and Cortical Default Mode networks demonstrated the strongest interaction effect (F = 10.43, FDR-corrected p = 0.0129), reflecting progressively divergent trajectories between groups across visits (Figure 1A). Specifically, Visual–Default connectivity increased over time in SCA7 but decreased in healthy controls, resulting in significantly stronger coupling in SCA7 at Visit 3. A second interaction effect was observed between the Cortical Somatomotor and Cerebellar Dorsal Attention networks (F = 8.04, FDR-corrected p = 0.0389), driven by a distinct longitudinal trajectory in SCA7 characterized by an increase from Visit 1 to Visit 2 followed by a partial return toward baseline levels (Figure 1B). These interaction effects are summarized in Figure 1C and indicate that longitudinal alterations in SCA7 preferentially involve interactions among Visual, Default Mode, Somatomotor, and cerebellar attention systems. Consistent with the significant Group × Visit interaction observed for the Cortical Somatomotor–Cerebellar Dorsal Attention network pair, post hoc within-group comparisons revealed a significant increase in connectivity from Visit 1 to Visit 2 in participants with SCA7 (Estimate = -0.2676, FDR-corrected p = 0.0062; Figure 1B). No other within-group longitudinal comparisons survived FDR correction.

**Figure 1.**
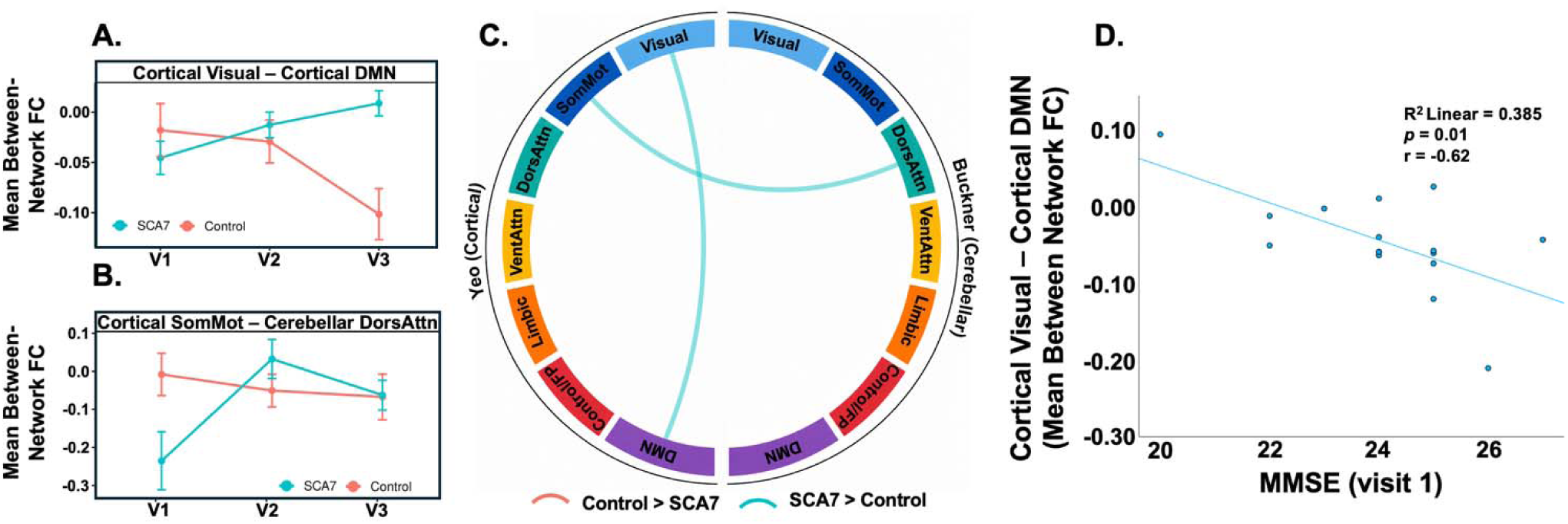
Longitudinal alterations in network-to-network functional connectivity and their association with cognitive performance in SCA7. (A) Mean functional connectivity between the Cortical Visual and Cortical Default Mode networks across the three study visits in healthy controls and participants with SCA7. (B) Mean functional connectivity between the Cortical Somatomotor and Cerebellar Dorsal Attention networks across visits. Error bars represent standard errors of the mean (SEM). (C Chord diagram illustrating network pairs exhibiting significant Group × Visit interaction effects. Network colors correspond to the seven cortical Yeo networks and seven cerebellar Buckner networks. (D) Scatterplot showing the association between Cortical Visual–Default Mode functional connectivity and baseline MMSE scores in the SCA7 group. The solid line represents the linear regression fit.

The clinical relevance of the Visual–Default interaction was further explored through correlation analyses. Visual–Default connectivity was negatively associated with baseline MMSE scores within the SCA7 group (r = -0.62, p = 0.01; Figure 1D), indicating that stronger coupling between these systems was associated with poorer cognitive performance. Together, these findings establish a relationship between altered Visual–Default connectivity and cognitive status in SCA7.

To further characterize the longitudinal interaction effects, between-group comparisons were examined at each visit to determine when these networks abnormalities emerged. At Visit 1, two between-group differences survived FDR correction (Figure 2A–B). Connectivity between the Cortical Dorsal Attention and Cerebellar Dorsal Attention networks was significantly lower in SCA7 than in healthy controls (Estimate = 0.2500, FDR-corrected p = 0.0233). In contrast, connectivity between the Cortical Somatomotor and Cortical Default Mode networks was significantly higher in SCA7 relative to controls (Estimate = -0.0985, FDR-corrected p = 0.0233). These findings indicate that abnormalities in both cortico-cerebellar and cortical large-scale network communication are already detectable at baseline assessment. Notably, stronger connectivity between the Cortical Somatomotor and Cortical Default Mode networks was associated with greater disease severity, as reflected by a positive correlation with baseline SARA scores within the SCA7 group (r = 0.589, p = 0.016; Figure 2C).

**Figure 2.**
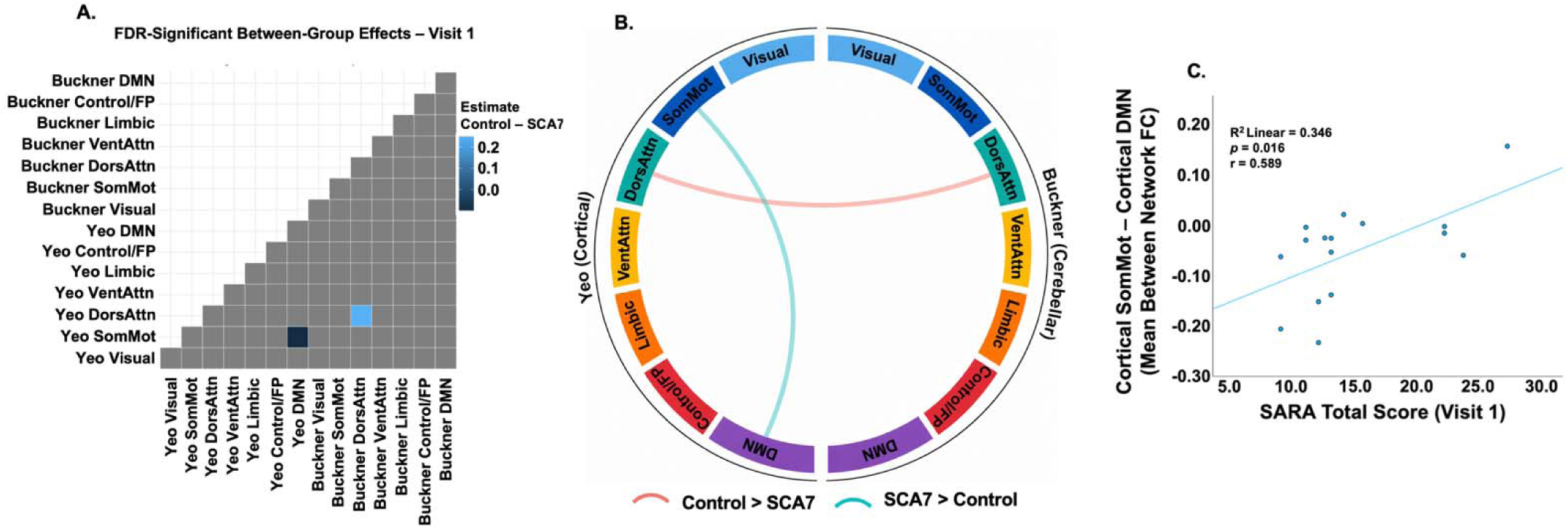
Baseline network-to-network functional connectivity differences and their association with ataxia severity in SCA7. (A) Matrix showing FDR-corrected between-group differences in network-to-network functional connectivity at Visit 1. Color intensity represents the estimated group difference (Control − SCA7). (B) Chord diagram illustrating significant between-group differences in network-to-network functional connectivity at Visit 1. Network colors correspond to the seven cortical Yeo networks and seven cerebellar Buckner networks. Red connections indicate greater connectivity in healthy controls, whereas teal connections indicate greater connectivity in participants with SCA7. (C) Scatterplot showing the association between Cortical Somatomotor–Default Mode functional connectivity and baseline SARA total scores within the SCA7 group. The solid line represents the linear regression fit.

By Visit 3, abnormalities became substantially more widespread. Among cortical–cortical interactions, participants with SCA7 demonstrated increased connectivity between the Cortical Visual and Cortical Default Mode networks (Estimate = -0.1104, FDR-corrected p = 0.0077), alongside reduced connectivity between the Cortical Visual and Cortical Dorsal Attention networks (Estimate = 0.1206, FDR-corrected p = 0.0119), the Cortical Visual and Cortical Somatomotor networks (Estimate = 0.1553, FDR-corrected p = 0.0119), and the Cortical Control and Cortical Default Mode networks (Estimate = 0.0797, FDR-corrected p = 0.0138) (Figure 3A, B). Notably, three of the four significant cortical abnormalities identified at Visit 3 involved the Visual network.

**Figure 3.**
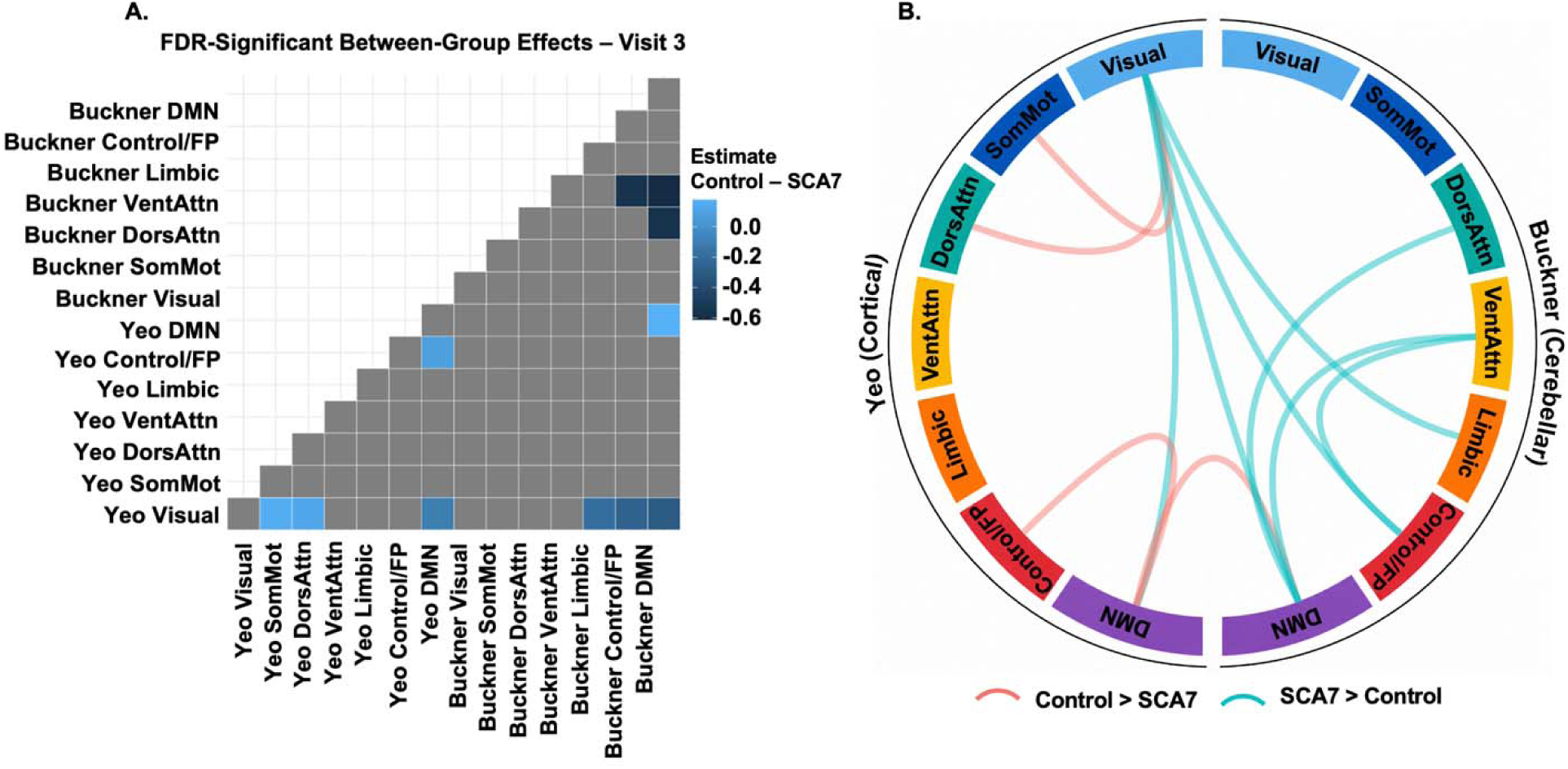
Between-group differences in network-to-network functional connectivity at Visit 3. (A) Matrix showing FDR-corrected between-group differences in network-to-network functional connectivity at Visit 3. Color intensity represents the estimated group difference (Control − SCA7). (B) Chord diagram illustrating significant cortical–cortical, cortico-cerebellar, and cerebellar–cerebellar network pairs identified at Visit 3. Network colors correspond to the seven cortical Yeo networks and seven cerebellar Buckner networks. Red connections indicate greater connectivity in healthy controls, whereas teal connections indicate greater connectivity in participants with SCA7.

Multiple cortico-cerebellar abnormalities also emerged at Visit 3. Relative to controls, SCA7 participants exhibited increased connectivity between the Cortical Visual network and the Cerebellar Default (Estimate = -0.3076, FDR-corrected p = 0.0119), Cerebellar Frontoparietal (Estimate = -0.2579, FDR-corrected p = 0.0188), and Cerebellar Limbic networks (Estimate = - 0.1927, FDR-corrected p = 0.0360). In contrast, connectivity between the Cortical Default and Cerebellar Default networks was reduced in SCA7 (Estimate = 0.1785, FDR-corrected p = 0.0119). This predominance of Visual-network abnormalities extended beyond cortical systems, as several cortico-cerebellar alterations involving the Visual network were also observed at Visit 3.

Cerebellar–cerebellar interactions were likewise altered at Visit 3. Compared with controls, SCA7 participants demonstrated increased connectivity between the Cerebellar Dorsal Attention and Cerebellar Default networks (Estimate = -0.5668, FDR-corrected p = 0.0277), between the Cerebellar Ventral Attention and Cerebellar Default networks (Estimate = -0.6123, FDR-corrected p = 0.0119), and between the Cerebellar Ventral Attention and Cerebellar Frontoparietal networks (Estimate = -0.5556, FDR-corrected p = 0.0154). A comprehensive summary of all FDR-significant network organization findings identified across global, network-specific, and between-system analyses is provided in Table 3.

**Table 3.**
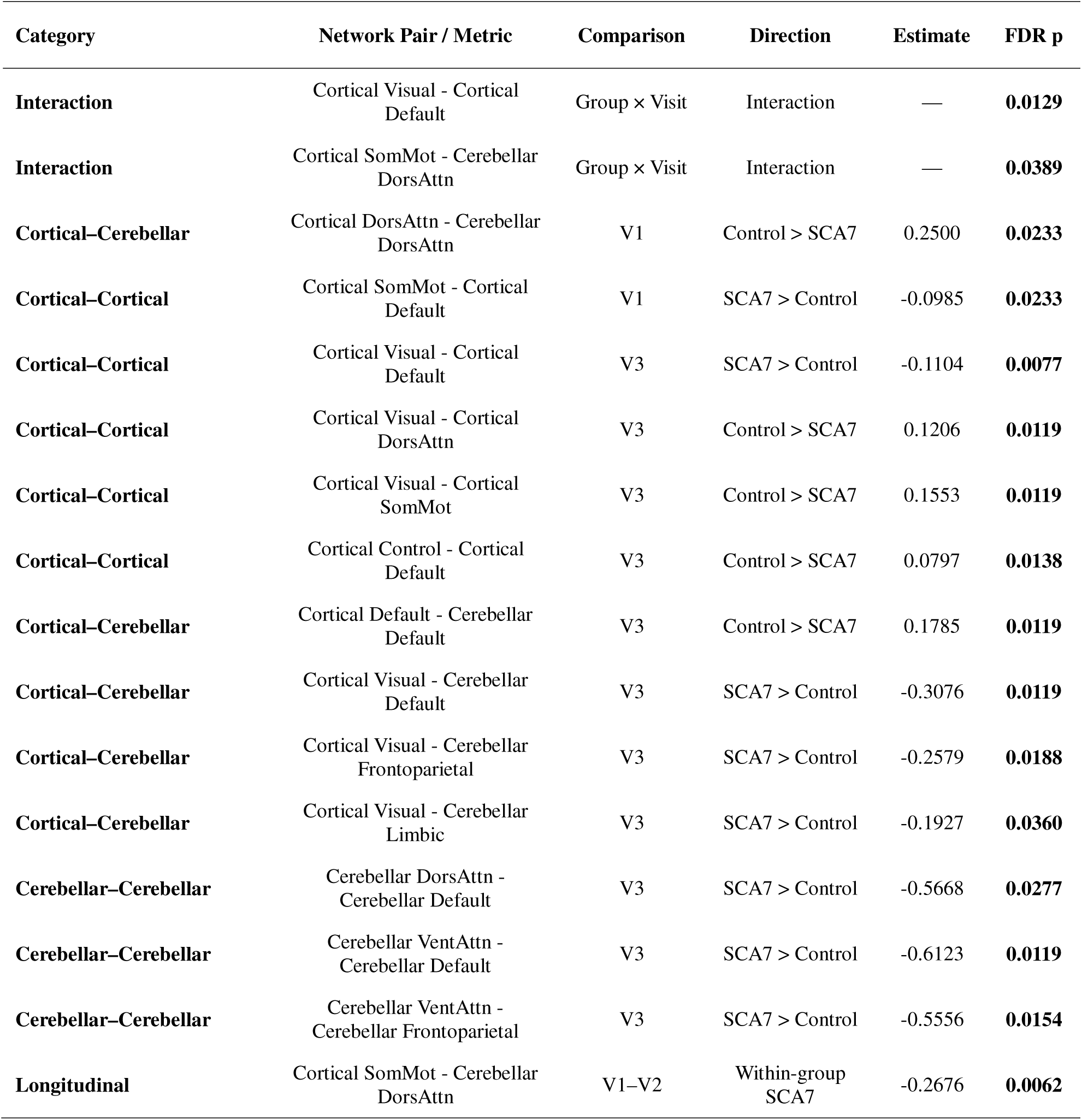
Summary of FDR-Significant Network Organization Findings in SCA7. DorsAttn = Dorsal Attention; SomMot = Somatomotor; Cont = Frontoparietal/Control network; SCA7, spinocerebellar ataxia type 7.

| Category | Network Pair / Metric | Comparison | Direction | Estimate | FDR p |
| --- | --- | --- | --- | --- | --- |
| **Interaction** | Cortical Visual - Cortical Default | Group × Visit | Interaction | — | **0.0129** |
| **Interaction** | Cortical SomMot - Cerebellar DorsAttn | Group × Visit | Interaction | — | **0.0389** |
| **Cortical–Cerebellar** | Cortical DorsAttn - Cerebellar DorsAttn | V1 | Control > SCA7 | 0.2500 | **0.0233** |
| **Cortical–Cortical** | Cortical SomMot - Cortical Default | V1 | SCA7 > Control | -0.0985 | **0.0233** |
| **Cortical–Cortical** | Cortical Visual - Cortical Default | V3 | SCA7 > Control | -0.1104 | **0.0077** |
| **Cortical–Cortical** | Cortical Visual - Cortical DorsAttn | V3 | Control > SCA7 | 0.1206 | **0.0119** |
| **Cortical–Cortical** | Cortical Visual - Cortical SomMot | V3 | Control > SCA7 | 0.1553 | **0.0119** |
| **Cortical–Cortical** | Cortical Control - Cortical Default | V3 | Control > SCA7 | 0.0797 | **0.0138** |
| **Cortical–Cerebellar** | Cortical Default - Cerebellar Default | V3 | Control > SCA7 | 0.1785 | **0.0119** |
| **Cortical–Cerebellar** | Cortical Visual - Cerebellar Default | V3 | SCA7 > Control | -0.3076 | **0.0119** |
| **Cortical–Cerebellar** | Cortical Visual - Cerebellar Frontoparietal | V3 | SCA7 > Control | -0.2579 | **0.0188** |
| **Cortical–Cerebellar** | Cortical Visual - Cerebellar Limbic | V3 | SCA7 > Control | -0.1927 | **0.0360** |
| **Cerebellar–Cerebellar** | Cerebellar DorsAttn - Cerebellar Default | V3 | SCA7 > Control | -0.5668 | **0.0277** |
| **Cerebellar–Cerebellar** | Cerebellar VentAttn - Cerebellar Default | V3 | SCA7 > Control | -0.6123 | **0.0119** |
| **Cerebellar–Cerebellar** | Cerebellar VentAttn - Cerebellar Frontoparietal | V3 | SCA7 > Control | -0.5556 | **0.0154** |
| **Longitudinal** | Cortical SomMot - Cerebellar DorsAttn | V1–V2 | Within-group SCA7 | -0.2676 | **0.0062** |

### 3.3 Classification Performance Using Functional Connectivity Features

Machine-learning models trained using resting-state functional connectivity features demonstrated high classification performance (Table 4). Among the three classifiers evaluated, Logistic Regression achieved the highest performance, with an accuracy of 96.4% and an F1 score of 0.969, followed closely by the Support Vector Machine (accuracy = 96.3%, F1 = 0.968). Random Forest showed comparatively lower classification performance. Given its superior performance, the Logistic Regression classifier was selected for further interpretation. LOOCV correctly classified 11 of 12 healthy controls and all 16 participants with SCA7, resulting in a single false-positive classification (Figure 4A).

**Figure 4.**
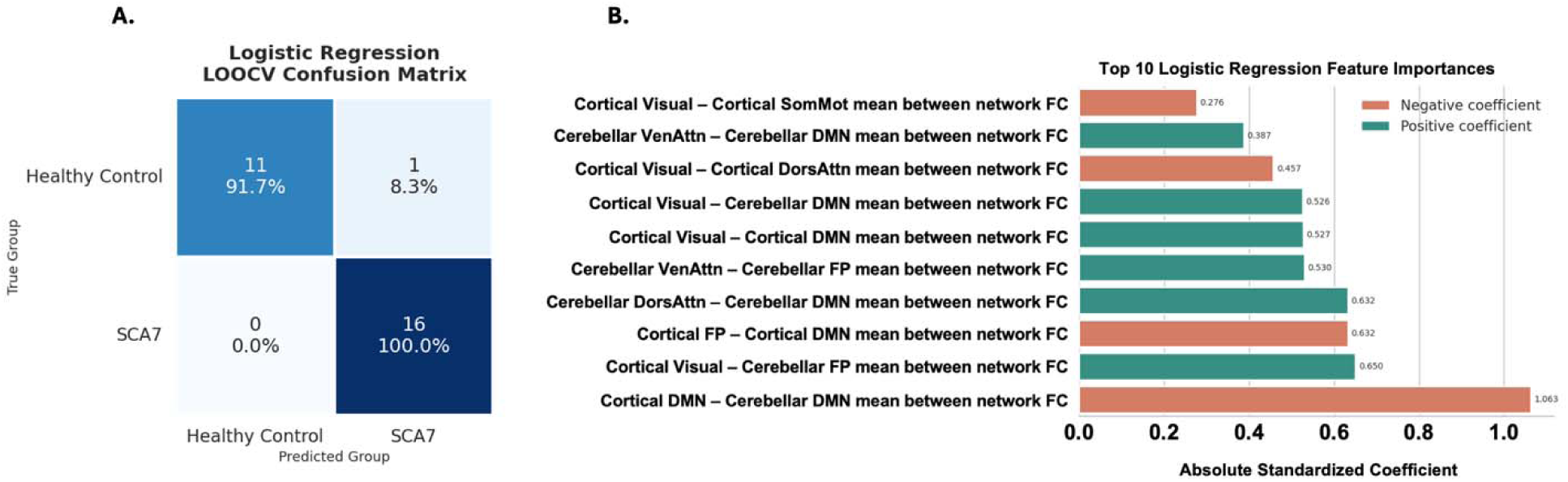
Classification performance and feature importance of the best-performing machine learning model.(A) Confusion matrix of the fMRI-only Logistic Regression classifier evaluated using Leave-One-Out Cross-Validation (LOOCV). (B) Feature importance rankings for the ten most influential functional connectivity features identified by the fMRI-only Logistic Regression model. Bar length represents the absolute standardized coefficient magnitude, and bar color indicates the direction of the coefficient (positive or negative).

**Table 4.** Performance of Machine Learning Models for Classifying SCA7 and Healthy Controls Using Functional Connectivity Features.

| Feature Set | Model | Accuracy (%) | F1 Score |
| --- | --- | --- | --- |
| fMRI (13 Features) | Logistic Regression | 96.4 | 0.969 |
|  | Random Forest | 82.1 | 0.857 |
|  | SVM | 96.3 | 0.968 |

Feature-importance analysis indicated that the most influential predictors primarily involved interactions among the Default Mode, Visual, and cerebellar networks (Figure 4B). The highest-ranked features included Cortical Default–Cerebellar Default, Cerebellar Dorsal Attention–Cerebellar Default, Cortical Visual–Cerebellar Frontoparietal, Cortical Visual–Cortical Default, and Cortical Visual–Cortical Somatomotor connectivity.

## 4. Discussion

This longitudinal resting-state fMRI study investigated large-scale functional network organization in SCA7 across three assessment time points. Several key findings emerged. First, participants with SCA7 exhibited alterations in global network organization characterized by reduced cortical within-network connectivity and increased between-network connectivity at baseline, together with increased global functional connectivity at the final assessment. Second, the Visual network emerged as the most consistently affected cortical system, demonstrating persistent abnormalities across visits and significant associations with disease severity. Third, network-level analyses identified altered interactions among Visual, Default Mode, Attention, Somatomotor, and cerebellar systems, with abnormalities becoming more widespread at later stages of follow-up and involving cortical, cortico-cerebellar, and cerebellar connections. Fourth, several connectivity measures were associated with clinical indices of motor and cognitive function, supporting their potential clinical relevance. Finally, machine-learning analyses demonstrated that resting-state functional connectivity features alone accurately distinguished individuals with SCA7 from healthy controls and outperformed models trained using behavioral measures. Collectively, these findings suggest that SCA7 is associated with progressive alterations in functional brain organization extending beyond the cerebellum and involving distributed cortical and cerebellar networks, highlighting the value of longitudinal resting-state fMRI for characterizing disease-related network reorganization.

Among all cortical systems examined, the Visual network emerged as the most consistently affected network. This finding is particularly noteworthy given that SCA7 is unique among the spinocerebellar ataxias in combining cerebellar degeneration with progressive retinal pathology and visual impairment(Campos-Romo et al., 2018; Miller et al., 2009). Previous structural and functional imaging studies have reported abnormalities involving visual pathways, occipital cortex, and visual-motor interactions, suggesting that visual dysfunction in SCA7 extends beyond retinal degeneration alone (Coarelli et al., 2024; Contreras et al., 2021a; Hernandez-Castillo, Galvez, et al., 2016b; Hernandez-Castillo, Vaca-Palomares, et al., 2016b; Horton et al., 2013b). The association between visual-network organization and SARA scores further supports the clinical relevance of these alterations and is consistent with prior studies linking imaging-derived measures to disease severity (Contreras et al., 2021a; Hernandez-Castillo, Vaca-Palomares, et al., 2016b). One notable aspect of the present findings is the remarkable convergence across analytical approaches. Visual-system abnormalities emerged in longitudinal network-to-network analyses, network-specific exploratory analyses, brain-behavior correlations, and machine-learning feature rankings. This consistency suggests that disruption of large-scale visual network organization may represent a central feature of functional brain alterations in SCA7.

Beyond the visual system, the present findings revealed abnormalities involving Default Mode, Attention, and Cerebellar networks, with network alterations becoming more extensive at later follow-up assessments. The cerebellum is increasingly recognized as a component of distributed networks supporting cognitive, attentional, and affective functions in addition to motor control (Koziol et al., 2012, 2014; Sokolov et al., 2017; Timmann et al., 2010). Accordingly, degeneration affecting cerebellar structures would be expected to influence communication across widespread cortical systems (Ackermann et al., 2007; Gellersen et al., 2017; Samson & Claassen, 2017; Sathyanesan et al., 2019). The observed cortico-cerebellar and cerebellar–cerebellar abnormalities are therefore consistent with previous evidence of widespread cerebro-cerebellar disruption in SCA7 (Chirino et al., 2018; Hernandez□Castillo et al., 2013; Lindsay & Storey, 2017). The repeated involvement of Default Mode network interactions and their association with cognitive performance further suggest that disease-related changes may extend beyond motor dysfunction and involve higher-order cognitive systems. This interpretation is consistent with growing evidence that cognitive impairment represents an important component of the SCA7 phenotype, although the specific cognitive implications of the observed connectivity alterations remain to be determined (Contreras et al., 2021a; Giocondo & Curcio, 2018; Petit et al., 2025). This broader pattern of distributed network involvement was also reflected in the complementary exploratory analyses presented in the Supplementary Material. Although these analyses did not demonstrate consistent longitudinal changes across visits and were therefore reported separately, they provided additional insight into large-scale network organization. Specifically, they suggested reduced cortical within-network connectivity together with greater between-network integration at baseline, consistent with diminished functional specialization. Network-specific analyses further demonstrated persistent disruption of the Visual network, reflected by reduced within-network connectivity across visits and altered network segregation, reinforcing the prominent role of visual-system dysfunction in SCA7.

An important aspect of the present findings is that several connectivity abnormalities were associated with clinical measures of motor and cognitive function. Previous neuroimaging studies in SCA7 have demonstrated relationships between disease severity and structural measures of gray matter, white matter, and spinal cord integrity (Contreras et al., 2021b; Hernandez-Castillo et al., 2021; Hernandez-Castillo, Galvez, et al., 2016b; Hernandez-Castillo, Vaca-Palomares, et al., 2016b). The current results extend this literature by suggesting that alterations in large-scale functional network organization may also reflect clinically meaningful aspects of disease burden. Rather than representing isolated imaging abnormalities, network-level connectivity changes appear to capture dimensions of disease expression that are relevant to both motor and cognitive outcomes.

The machine-learning analyses further support the relevance of large-scale functional connectivity alterations in SCA7. These findings are consistent with previous work demonstrating that whole-brain functional connectivity can distinguish individuals with SCA7 from healthy controls with high accuracy (Hernandez-Castillo et al., 2014). Particularly noteworthy is that several of the highest-ranking features in the present model involved Visual, Default Mode, and cerebellar systems, which were also implicated by the group-level analyses. This convergence across independent analytical approaches strengthens confidence that the identified network abnormalities reflect meaningful disease-related characteristics rather than statistical artifacts. Nevertheless, the classification findings should be interpreted cautiously given the modest sample size and the absence of external validation. Because both feature selection and model training were performed in a relatively small cohort, the reported classification performance should be regarded as preliminary and may not generalize to independent samples. Future multicenter studies will be necessary to determine the robustness and potential translational utility of these models.

Several strengths and limitations should be considered when interpreting the present findings. A major strength of this study is its longitudinal design, which allowed the characterization of functional network alterations across multiple time points in a relatively rare neurodegenerative disorder. In addition, the use of complementary analytical approaches, including global network metrics, network-specific analyses, clinical association testing, and machine-learning classification, provided a comprehensive assessment of large-scale functional organization in SCA7. However, the modest sample size may have limited statistical power to detect more subtle longitudinal effects. Clinical data were also unavailable for some participants and time points, restricting the evaluation of longitudinal brain-behavior relationships. Furthermore, direct measures of retinal structure and visual function were not available, limiting interpretation of the visual-network findings. Finally, the Buckner cerebellar parcellation represents each cerebellar network as a single node, necessitating the exclusion of cerebellar networks from certain segregation analyses and limiting the assessment of within-network cerebellar organization.

The present findings support a network-based view of SCA7, demonstrating that the disorder is associated with widespread alterations in functional communication across cortical and cerebellar systems rather than isolated cerebellar pathology alone. In particular, the prominent involvement of visual and cerebro-cerebellar networks suggests that disease-related changes extend across multiple interconnected systems that support sensory, motor, and cognitive functions. By characterizing these alterations longitudinally, this study provides further insight into the large-scale functional reorganization associated with SCA7 and highlights the value of network-level approaches for understanding the neural mechanisms underlying disease progression.

## Data availability

Data will be made available upon request.

## Funding

This work was supported by the following grants and granting agencies: the Natural Sciences and Engineering Research Council of Canada (NSERC) grant RGPIN-2022-03368; Canada Research Chair grant CRC-2020-00079 to CRH-C; Universidad Nacional Autonoma de Mexico DGAPA-PAPIIT IN208625 to JFR.

## Competing interests

The authors report no competing interests.

## Supplementary material

Supplementary material is available as a separate file.

## Notes

### Competing Interest Statement

The authors have declared no competing interest.

### Summary of Updates

We recently received the approval email for our bioRxiv submission and would like to request a change to the manuscript title. Current title: Longitudinal Reorganization of Cortical and Cerebellar Functional Networks in Spinocerebellar Ataxia Type 7 New title: Longitudinal Reorganization of Large-Scale Functional Networks in SCA7

