## supplementary materials for "Longitudinal Reorganization of Large-Scale Functional Networks in SCA7"

**Exploratory Analyses of Global Functional Connectivity and Network Organization**

To provide a broader characterization of large-scale functional brain organization, complementary exploratory analyses of global functional connectivity and network organization were performed. These analyses are presented in the Supplementary Material because they demonstrated limited longitudinal effects and primarily served to provide additional context for the primary network-to-network findings reported in the main manuscript.

***Global Functional Connectivity***

Global functional connectivity was quantified as a summary measure of large-scale functional organization. Using the seven cortical functional systems defined by the Yeo-7 atlas and the seven cerebellar functional systems defined by the Buckner-7 atlas, pairwise Pearson correlation coefficients were calculated between all system time series for each participant and visit, yielding a 14 × 14 functional connectivity matrix. Correlation coefficients were transformed to Fisher z-scores prior to analysis. Mean whole-brain functional connectivity was calculated as the average of all unique pairwise connectivity values within each matrix, resulting in a single summary measure of global resting-state functional connectivity for each scan.

***Network Organization Analysis***

Large-scale network organization was examined using measures of network segregation and between-system integration (Chan et al., 2014). Analyses were performed using the Schaefer-200 cortical parcellation, with cortical parcels assigned to the seven canonical Yeo functional networks (Visual, Somatomotor, Dorsal Attention, Salience/Ventral Attention, Limbic, Frontoparietal/Control, and Default Mode). For analyses involving cortico-cerebellar and cerebellar interactions, the seven cerebellar systems defined by the Buckner-7 atlas were additionally incorporated.

Within-network functional connectivity was calculated as the mean Fisher z-transformed connectivity among parcels belonging to the same network, whereas between-network functional connectivity was calculated as the mean connectivity between parcels belonging to different networks. Network segregation was quantified as:

$$System Segregation= \frac{(Z̄within - Z̄between)}{Z̄within}$$

where Z̄within represents the mean Fisher z-transformed connectivity among parcels within the same network and Z̄between represents the mean Fisher z-transformed connectivity between parcels belonging to different networks. Higher segregation values indicate greater functional specialization and stronger separation between a network and the remainder of the brain, whereas lower values indicate reduced network differentiation and greater cross-network integration.

Network segregation was evaluated at both global and network-specific levels. Global cortical segregation was computed using mean within-network and between-network connectivity values aggregated across all cortical networks. Network-specific analyses were subsequently performed for each cortical network individually, including assessments of within-network connectivity, between-network connectivity, and network segregation.

***Group Differences in Global Functional Connectivity Emerged at the Final Follow-Up Visit***

Mean whole-matrix functional connectivity values for each group and visit are presented in Figure 1. The linear mixed-effects model did not reveal a significant Group × Visit interaction (F(2,52) = 2.11, p = 0.132). Nevertheless, post-hoc analyses demonstrated significantly higher mean whole-matrix functional connectivity in the SCA7 group compared with healthy controls at Visit 3 (Estimate = -0.0538, t = -2.430, p = 0.018), whereas no significant between-group differences were observed at Visits 1 or 2.

Within-group analyses indicated stable global functional connectivity across visits in healthy controls. In contrast, participants with SCA7 exhibited a significant increase in mean whole-matrix functional connectivity between Visit 1 and Visit 3 (Estimate = -0.0384, t = -2.542, p = 0.037). A similar increase was observed between Visit 2 and Visit 3, although this comparison did not reach statistical significance (Estimate = -0.0354, t = -2.341, p = 0.059).

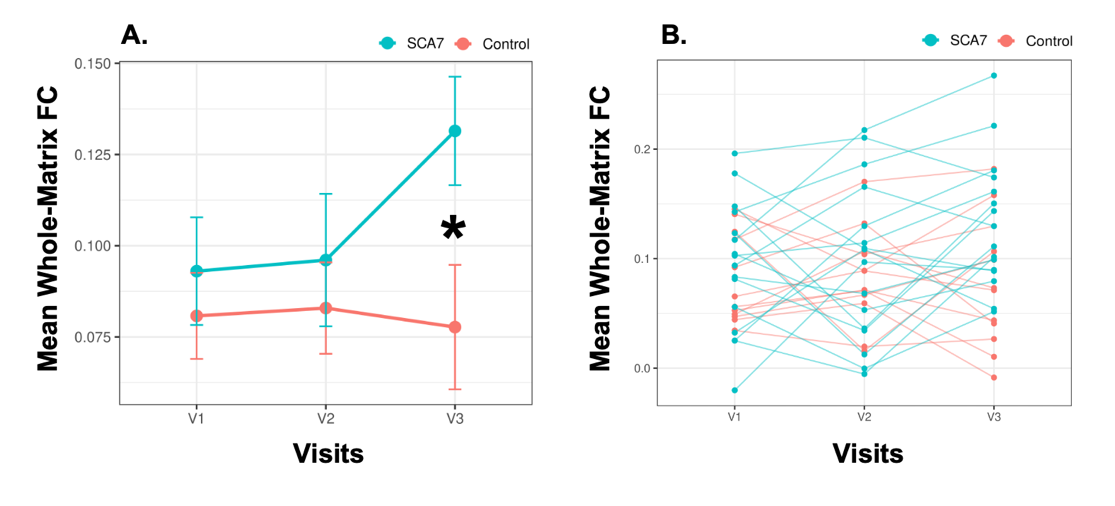
Overall, these findings suggest that group differences in global functional connectivity became more pronounced at the final follow-up visit, primarily driven by increasing connectivity within the SCA7 group over time.

***Figure1.*** Longitudinal changes in mean whole-matrix functional connectivity in healthy controls and participants with SCA7. (A) Group-level mean whole-matrix functional connectivity (Fisher z-transformed) across the three study visits. Error bars represent standard errors of the mean (SEM). Participants with SCA7 exhibited significantly higher mean whole-matrix functional connectivity than healthy controls at Visit 3 (p = 0.018), indicated by the asterisk. No significant between-group differences were observed at Visits 1 or 2. (B) Individual participant trajectories across visits. Healthy controls demonstrated stable mean whole-matrix functional connectivity across all visits, whereas participants with SCA7 exhibited a significant increase in connectivity between Visit 1 and Visit 3 (p = 0.037), with a trend-level increase observed between Visit 2 and Visit 3 (p = 0.059). Although the overall Group × Visit interaction was not statistically significant (F(2,52) = 2.11, p = 0.132), the longitudinal trajectories suggest increasing divergence in global functional connectivity over time, driven primarily by elevated connectivity in the SCA7 group at the final follow-up visit.

***SCA7 Exhibited Reduced Within-Network Coherence and Increased Between-Network Integration at Baseline***

Global cortical system segregation, mean within-network cortical functional connectivity, and mean between-network cortical functional connectivity are shown in Figure 2. Linear mixed-effects analyses revealed no significant Group × Visit interactions for global cortical segregation (FDR-corrected p = 0.743), mean within-network cortical functional connectivity (FDR-corrected p = 0.743), or mean between-network cortical functional connectivity (FDR-corrected p = 0.743).

Despite the absence of significant longitudinal effects, post-hoc analyses identified two between-group differences that survived FDR correction at Visit 1. Participants with SCA7 exhibited significantly lower mean within-network cortical functional connectivity relative to healthy controls (Estimate = 0.0587, FDR-corrected p = 0.024), indicating reduced coherence within canonical cortical systems. In addition, SCA7 participants demonstrated significantly less negative between-network cortical functional connectivity compared with controls (Estimate = -0.0215, FDR-corrected p = 0.040), suggesting greater functional integration and reduced separation between large-scale cortical networks.

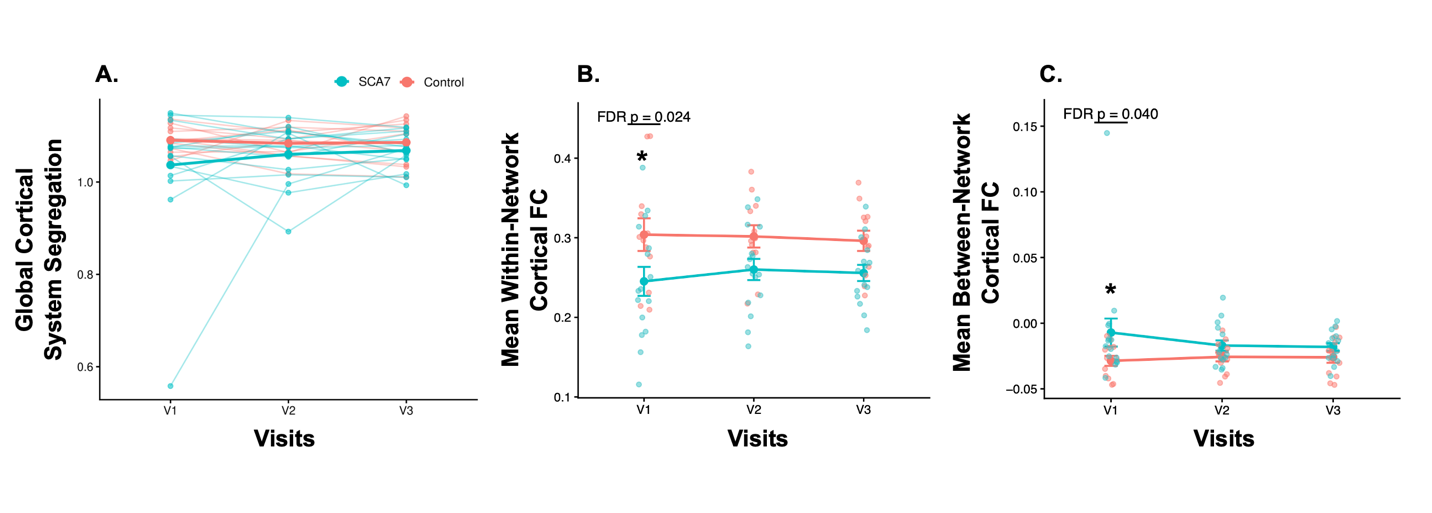
Although the global cortical segregation index did not survive FDR correction, both contributing components showed alterations in the expected direction. Specifically, reduced within-network connectivity together with increased between-network connectivity was consistent with a shift toward reduced systems-level specialization in SCA7. No within-group longitudinal comparison survived FDR correction for any of the three global network-organization metrics.

***Figure 2.*** Global cortical network segregation and integration metrics in healthy controls and participants with SCA7. (A) Global cortical system segregation across the three study visits. System segregation was calculated using cortical parcels from the Schaefer-200 atlas grouped according to the Yeo 7-network framework. Individual participant trajectories are shown together with group means. No significant Group × Visit interaction or within-group longitudinal effects survived FDR correction. (B) Mean within-network cortical functional connectivity across visits. Participants with SCA7 exhibited significantly lower within-network cortical functional connectivity than healthy controls at Visit 1 (FDR-corrected p = 0.024), indicating reduced coherence within canonical cortical systems. (C) Mean between-network cortical functional connectivity across visits. Participants with SCA7 exhibited significantly higher (i.e., less negative) between-network cortical functional connectivity than healthy controls at Visit 1 (FDR-corrected p = 0.040), indicating reduced separation and greater integration between large-scale cortical networks. Error bars represent standard errors of the mean (SEM). Asterisks denote FDR-significant between-group differences. No within-group longitudinal comparisons survived FDR correction for any of the three metrics.

***Persistent Visual-Network Abnormalities and Their Clinical Associations in SCA7***

Network-specific cortical analyses were performed using the Schaefer-200 cortical parcellation grouped according to the Yeo seven-network framework. No Group × Visit interaction survived FDR correction for any cortical network or metric.

As illustrated in Figure 3A, the most consistent network-specific finding involved the Visual network, which demonstrated significantly lower within-network functional connectivity in participants with SCA7 relative to healthy controls at Visit 1 (Estimate = 0.1501, FDR-corrected p = 0.0256), Visit 2 (Estimate = 0.1500, FDR-corrected p = 0.0258), and Visit 3 (Estimate = 0.1435, FDR-corrected p = 0.0375). Reduced within-network connectivity was also observed within the Frontoparietal/Control network at Visit 2 (Estimate = 0.0794, FDR-corrected p = 0.0323). No within-group longitudinal comparison survived FDR correction for any cortical network or metric.

To assess the clinical relevance of these findings, correlation analyses were performed between significant network measures and clinical variables. As shown in Figure 3B, Visual-network within-network functional connectivity was negatively associated with baseline SARA total scores within the SCA7 group (r = -0.687, p = 0.003), indicating that lower Visual-network coherence was associated with greater ataxia severity. Collectively, these findings identify the Visual system as the most consistently affected cortical network in SCA7 and suggest that disruption of Visual-network organization is related to clinical disease burden.

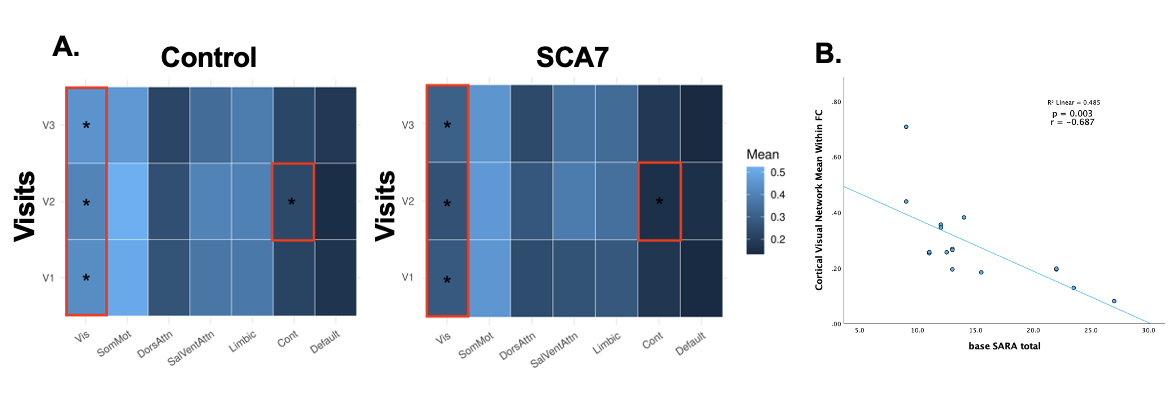
***Figure 3.*** Persistent visual-network abnormalities and their association with ataxia severity in SCA7. (A) Heatmaps depicting mean within-network functional connectivity (Fisher z-transformed) for each cortical Yeo network across visits in healthy controls and participants with SCA7. Asterisks denote networks demonstrating significant between-group differences following FDR correction. The Visual network exhibited significantly reduced within-network functional connectivity in SCA7 relative to controls at all three visits (Visit 1: FDR-corrected p = 0.0256; Visit 2: FDR-corrected p = 0.0258; Visit 3: FDR-corrected p = 0.0375). Reduced within-network connectivity was also observed in the Frontoparietal/Control network at Visit 2 (FDR-corrected p = 0.0323). Red boxes highlight networks exhibiting FDR-significant between-group differences. (B) Association between Visual-network within-network functional connectivity and baseline SARA total scores within the SCA7 group. Lower Visual-network connectivity was associated with greater ataxia severity (r = -0.687, p = 0.003). Together, these findings identify the Visual system as the most consistently disrupted cortical network in SCA7 and demonstrate a significant relationship between Visual-network integrity and clinical disease severity.
